# Bayesian Energy Landscape Tilting: Towards Concordant Models of Molecular Ensembles

**DOI:** 10.1101/002048

**Authors:** Kyle A. Beauchamp, Vijay S. Pande, Rhiju Das

**Keywords:** Molecular Dynamics, NMR, Conformational Ensembles, Bayesian Statistics

## Abstract

Predicting biological structure has remained challenging for systems such as disordered proteins that take on myriad conformations. Hybrid simulation/experiment strategies have been undermined by difficulties in evaluating errors from computational model inaccuracies and data uncertainties. Building on recent proposals from maximum entropy theory and nonequilibrium thermodynamics, we address these issues through a Bayesian Energy Landscape Tilting (BELT) scheme for computing Bayesian “hyperensembles” over conformational ensembles. BELT uses Markov chain Monte Carlo to directly sample maximum-entropy conformational ensembles consistent with a set of input experimental observables. To test this framework, we apply BELT to model trialanine, starting from disagreeing simulations with the force fields ff96, ff99, ff99sbnmr-ildn, CHARMM27, and OPLS-AA. BELT incorporation of limited chemical shift and ^3^*J* measurements gives convergent values of the peptide’s *α*, *β*, and *PP*_*II*_ conformational populations in all cases. As a test of predictive power, all five BELT hyperensembles recover set-aside measurements not used in the fitting and report accurate errors, even when starting from highly inaccurate simulations. BELT’s principled fxramework thus enables practical predictions for complex biomolecular systems from discordant simulations and sparse data.

## Introduction

The past forty years have seen the experimental determination of “ground-state” structures for countless biological macromolecules (1). Modern biology, however, presents many systems that do not fit a single-structure paradigm. “Excited” conformational states of nucleic acids (2), natively disordered proteins (3), and protein folding intermediates (4) are all poorly described by single conformation models. For such systems, models of conformational ensembles are required to understand and to predict structural and equilibrium properties.

A growing body of research has sought to characterize structural ensembles. Much of this work has focused on incorporating dynamical information during NMR structure determination (5, 6) or the extraction of multiple conformers from X-ray diffraction data (7, 8). While these techniques are powerful, they share difficulties in data collection, the unified treatment of heterogeneous experimental data, and data sparseness relative to the number of degrees of freedom. In particular, conformational ensemble modeling requires the estimation of not just a single structure, but a collection of structures and their associated equilibrium populations. This highly under-determined problem involves the simultaneous estimation of approximately 3 × *N* × *m* parameters, where m is the number of states in the ensemble and *N* is the number of atoms in the molecule. Estimating uncertainties of these ensembles further amplifies this challenge. Inference in this regime necessarily requires more information, which in principle can be attained by combining measurements with simulations that leverage prior physical understanding encoded in atomistic force fields.

Despite recent advances in force field development (9, 10), simulation benchmark studies have demonstrated continuing inaccuracies in molecular dynamics (MD) force fields (11). Force field modifications based on direct fitting to NMR measurements have also been demonstrated (12, 13, 14), but such work has optimized only a small fraction of the required force field parameters. Thus, simulations are often unable to recapitulate *ab initio* the wide variety of measurements available on molecular systems. This inaccuracy poses a challenge when one desires atomic-scale models that are both consistent with presently available measurements and predictive of those yet to be measured.

Here, we introduce a practical statistical approach to modeling solution ensembles of biological macromolecules. The algorithm, Bayesian Energy Landscape Tilting (BELT), uses solution experiments to reweight an ensemble of atomistic models predicted (perhaps inaccurately) by molecular dynamics. BELT generalizes a recently proposed maximum entropy method (15) to the practical scenario in which the experimental measurements and their estimated relationships to atomic conformation carry error. In particular, BELT leverages Markov Chain Monte Carlo (16) to transform experimental ambiguity into error bars on arbitrary structural features. The final output of BELT modeling is a hyperensemble, or an “ensemble of ensembles”, which we show is closely connected to a generalized ensemble theory proposed by Crooks (17). This hyperensemble is a collection of statistical samples, each of which is itself a conformational ensemble that corresponds to a maximum-entropy solution associated with a particular set of experimental observables.

The necessity and utility of this approach can be illustrated with a simple example with one experimental observable. Most previous methods have focused on obtaining estimates of a single best-fit conformational ensemble (15, 18, 19). However, ambiguous experimental data often disallow such a point-estimate of the conformational ensemble. For example, we plot one measured (19) value of ^3^*J*(*H*_*N*_*H*^*α*^) in the context of the Karplus (20) equation relating *φ* to ^3^*J*(*H*_*N*_*H*^*α*^) (Fig. 1a). The measured coupling corresponds to four different values of *ϕ*, precluding description by a single point estimate of *ϕ*, much less a single estimate of the distribution of *ϕ*. Many different ensembles are consistent with the measurement (Figs. 1b,c), leading to nearly completely loss of predictive power. A molecular dynamics simulation can establish a prior estimate for the ensemble (Fig. 1d), but may disagree with the observed data beyond measurement error (Fig. 1e). In this case, how to compute a statistical collection of ensembles that leverages both the simulation and the data has not been obvious; for example, prior Bayesian approaches return uncertainties assuming single conformations, not full ensembles (21). The BELT approach described herein (blue traces in Fig. 1d,e) offers a practical recipe for describing such a hyperensemble, for computing the hyperensemble’s predictions for new experimental observables not used in the modeling, and for giving rigorous error estimates on these predictions.

**Figure 1.**
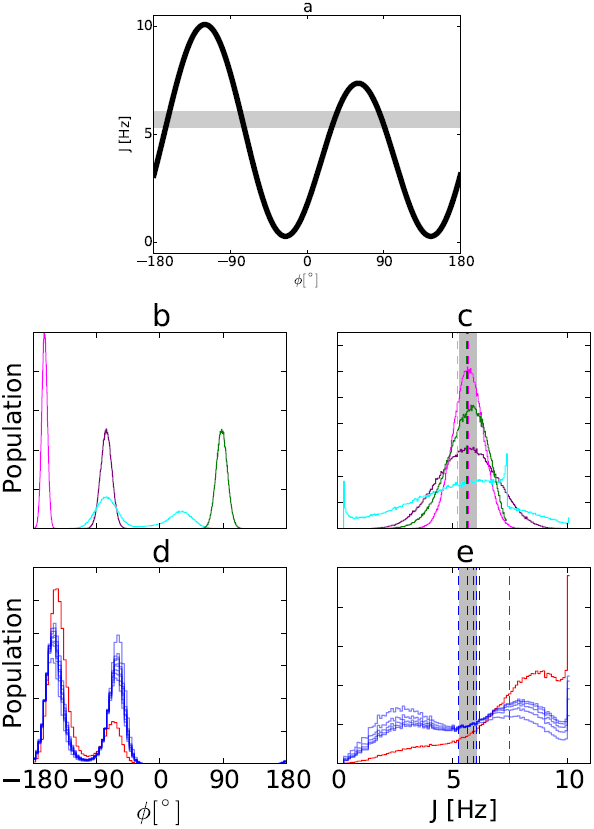
(a). The Karplus equation connecting the backbone torsion *ϕ* to ^3^*J*(*H*_*N*_*H*^*α*^). The measured value of ^3^*J*(*H*_*N*_*H*^*α*^) is shaded gray and is consistent with multiple values of *ϕ*. (b, c). Histograms of four chemically unrealistic ensembles that recapitulate the measured (gray) value of ^3^*J*(*H*_*N*_*H*^*α*^). Each histogram is represented in both the backbone torsion *ϕ* (b) and the projection (via Karplus equation) onto ^3^*J*(*H*_*N*_*H*^*α*^) (c). Dashed vertical bars represent the average ^3^*J*(*H*_*N*_*H*^*α*^) for each corresponding ensemble. (d, e). The molecular dynamics (ff99) ensemble (red) is inconsistent with the measured ^3^*J*(*H*_*N*_*H*^*α*^). Four samples (blue) from the BELT hyperensemble show good agreement with measured values of ^3^*J*(*H*_*N*_*H*^*α*^). For this figure only, the uncertainty (*σ*) on ^3^*J*(*H*_*N*_*H*^*α*^) was increased 2.5-fold to better illustrate differences. Density spikes in (c, e) correspond to values where 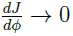.

After laying out the theoretical framework for BELT, this study presents in-depth tests based on assessing the convergence of ensembles constructed from force fields with radically different properties. We investigated the conformational propensities of trialanine using NMR measurements (19) and MD simulations performed in five different force fields. The small size of this model system enabled assessment of BELT without complications from incomplete sampling. At the same time, trialanine populates multiple conformational states and allows incisive tests of ensemble modeling. Although the raw simulations show wide variations in their conformational preferences, BELT corrects force field errors to provide concordant estimates of the *α*, *β*, and *PP*_*II*_ populations. The ability to correct the biases of diverse forcefields provides a stringent test of the proposed calculation scheme for connecting simulation and equilibrium measurements.

## Theory: Bayesian Energy Landscape Tilting

### Model Inputs

To model an ensemble using BELT requires three components. First, we need conformations *x*_*j*_ (*j* = 1,…,*m*) sampled from the equilibrium distribution of some physically realistic model. This model will serve as a prior on structural properties; in the absence of experimental data, the BELT model inherits the properties of the conformations *x*_*j*_. In the present work, such conformations will be generated from molecular dynamics simulations. Second, we require equilibrium experimental measurements *F*_*i*_ (*i* = 1,…,*n*) and their associated uncertainties *σ*_*i*_ (*i* = 1, …, *n*). Third, it is necessary to have a direct connection between simulation and experiment. This connection is achieved by predicting each experimental observable at each conformation: *f*_*i*_(*x*_*j*_) is the predicted value of experiment *i* at conformation *x*_*j*_.

### Reweighting

The next step in constructing an ensemble is to calculate the population of each conformation. Inspired by a previous method for restraining simulations (15) (see Appx. S1), we reweight individual conformations by a biasing potential that is a linear combination of the predicted observables:

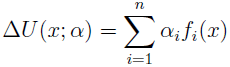

In Δ*U*(*x*; *α*), the parameters *α*_*i*_ determine how strongly each experiment contributes to the biasing potential. As shown previously (15), such a linear biasing potential gives a maximum entropy ensemble for some set of experimental observations. The BELT strategy is to look beyond the single best such ensemble so as to estimate the uncertainty in the ensemble modeling. BELT instead samples over a distribution of such maximum entropy ensembles each parametrized by *α*_*i*_. This approach is connected (see Appx. S1) to recent work by Crooks that proposed an entropic prior for modeling hyperensembles in general physical problems.

The end result is a collection of ‘landscape-tilted’ ensembles (Fig. 1e). That is, each conformational ensemble is a perturbed version of the initial molecular dynamics ensemble but reweighted (see Appx. S2) according to energetic perturbations that are linear in the experimental observables *f*_*i*_(*x*):

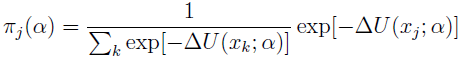

With the equilibrium populations, we can calculate the equilibrium expectations of an arbitrary observable *h*(*x*):

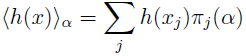

In the above bracket notation, 〈*h*(*x*)〉_*a*_ is the ensemble average of *h*(*x*) in an ensemble that is perturbed by a biasing potential Δ*U*(*x*; *α*). At this point, the determination of the parameters *α*_*i*_ has not yet been discussed. The key idea, however, is that the *α* reweighted ensemble 〈〉_*α*_ should recapitulate the experimental measurements:

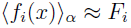

Forcing this to be an exact equality recovers previous results (15) that can be derived from maximum entropy considerations (Appx. S1); here, however, we take into account the experimental uncertainties associated with each *F*_*i*_.

## Determining *α*

A Bayesian framework enables determination of the coefficients *α* used in the biasing potential. An alternative derivation using the Crooks hyperensemble formalism (17) is given in Appx. S1. BELT assumes that, given the correct choice of α, the predicted observables *f*_*i*_(*x*) provide unbiased (but noisy) predictions of the measurements *F*_*i*_. This recipe assumes independence (see Appx. S3) and the following conditional probabilities:

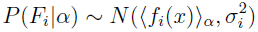

In the above equation, *N*(.,.) refers to a normal distribution with specified mean and variance. For the current work, we model *σ*_*i*_ as the uncertainty associated with predicting chemical shifts and scalar couplings from structures; this error is quantified by the RMS uncertainty estimated during the parameterization of chemical shift and scalar coupling models. Using Bayes’ Theorem, we can calculate the posterior distribution of *α*:

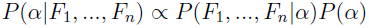

Now we let *LP*(*α*) denote the log posterior of α and simplify, dropping terms that are independent of *α*:

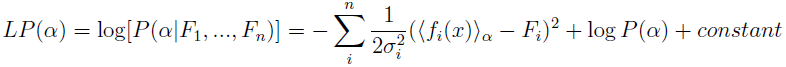

Note the simple form of the log posterior. The first term (i.e. the log likelihood) measures the *χ*^2^ agreement between the reweighted ensemble and measurements. The second term is the log of the prior distribution on *α*.

In the present work, we evaluate three different choices of prior (Appx. S4), finding similar results for each. The first is the maximum entropy (maxent) prior, which penalizes ensembles as they deviate from the raw simulation results:

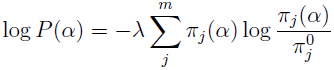

In the previous expression, 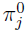 refers to the populations of an unweighted ensemble, which are typically 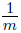, while λ is a hyperparameter that controls the strength of the prior. We also consider using a Dirichlet prior, which is functionally similar to the maxent prior (Appx. S4):

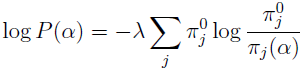

The third prior we consider is a multivariate normal prior, where *α* ~ *N*(0, Σ). The value of Σ is given by Σ_*ij*_ = λ*Cov*(*f*_*i*_(*x*), *f*_*j*_(*x*)), as derived in Appx. S4.

Each of these priors can be used to achieve regularization, which is a powerful technique to reduce overfitting (22). Large values of λ favor the raw simulation results (i.e. uniform conformational populations): 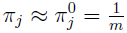. The value of λ can be chosen via cross-validation or other methods (see Appx. S5). When using the maxent prior in the limit of large λ and *σ* → 0, BELT recovers the hyperensemble picture of nonequilibrium statistical mechanics as developed (17) by Crooks (see Appx. S1). The Dirichlet and Normal priors do not share the same connection to the Crooks hyperensemble formalism; however, for normally distributed observables, all three priors will give identical results (23).

## MCMC Sampling of Structural Ensembles

As noted above, because ensemble inference often presents many plausible solutions (21, 24, 25), we avoid statistical methods that return a single solution (e.g. maximum likelihood or maximum entropy). We therefore use Markov chain Monte Carlo (MCMC), as implemented in PyMC (16), to sample the distribution of structural ensembles—one ensemble per sampled *α*—consistent with experiment. The result is an ensemble of ensembles—a statistical ensemble of conformational ensembles. Averaging all MCMC samples provides posterior mean estimates of arbitrary structural features or experimental observables. Similarly, examining the MCMC variances provides statistical uncertainties of equilibrium or structural features. A Bayesian bootstrapping procedure (26) can also be used to model the statistical uncertainty of the MD simulations (see Appx. S6).

## Methods

### Molecular Dynamics Simulations

Trialanine was simulated in the ff96 (27), ff99 (28), ff99sbnmr-ildn (29, 30), CHARMM27 (31, 32), and OPLS-AA (33) force fields, as previously reported (11). Simulations were performed using Gromacs 4.5 (34) and run at constant temperature (300 K) and pressure (1.01 atm). Each simulation was at least 225 ns long and used the TIP4P-EW water model (35). Conformations were stored every 1 ps.

### Chemical Shifts and Scalar Couplings

All NMR measurements in this work refer to experiments probing the central residue of trialanine (19). The experimental data were measured at pH 2, near the pKa of the carboxylate moiety of the C terminus, which would requires a constant pH simulation, rather than a fixed protonation state. Because such simulations are challenging with current force fields and simulation packages, we simulated the trialanine construct with charged termini— conditions in which the the force fields have been best calibrated and tested. We therefore focus our analysis on the central alanine residue, which should be most robust to pH dependent effects. Both pH differences and force field inaccuracies will lead to systematic differences between simulation and experiment; indeed, we assess whether BELT robustly corrects these deviations.

Chemical shifts (*H*, *H*^*α*^, *C*^*α*^, *C*^*β*^) for each frame were calculated using a weighted average of ShiftX2 (36), SPARTA+ (37), and PPM (38) predictions; uncertainties for each model were estimated using their reported RMS prediction errors. Overall uncertainties were estimated as 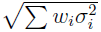, where 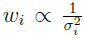 is the weight (Σ_*i*_*w*_*i*_ = 1) of each chemical shift model and *σ*_*i*_ is the uncertainty of each chemical shift model. The J couplings were calculated using the following Karplus relations: ^3^*J*(*H*^*N*^*C*′) (20), ^3^*J*(*H*^*N*^*H*^*α*^) (20), ^2^*J*(*NC*^*α*^) (19), ^3^*J*(*H*^*α*^*C*′) (39), ^1^*J*(*NC*^*α*^) (19), ^3^*J*(*H*^*N*^*C*^*β*^) (20). J coupling uncertainties were approximated as the RMS errors reported when fitting the Karplus coefficients.

We have divided the available experimental measurements into training and test sets, with the training set consisting of the ^3^*J*(*H*^*N*^*C*′), ^2^*J*(*NC*^*α*^), and ^3^*J*(*H*^N^*C*^*β*^) scalar couplings and the *C*^*α*^, *H*^*N*^, and *C*^*β*^ chemical shifts. The test set consists of ^3^*J*(*H*^*N*^*H*^*α*^), ^3^*J*(*H*^*α*^*C*′), ^1^*J*(*NC*^*α*^), and the *H*^*α*^ chemical shift. The division into training and test sets serves three purposes. First, it provides a test of overfitting. Second, it allows us to reduce the computational cost of BELT calculations. Third, it allows us to train on data that are approximately uncorrelated; BELT is best suited for working with uncorrelated data.

## BELT

All BELT calculations were performed using the FitEnsemble package (https://github.com/kyleabeauchamp/FitEnsemble). The online FitEnsemble tutorial demonstrates the use of BELT with a single experimental measurement (^3^*J*(*H*^*N*^*H*^*α*^)). Source code for calculations in this work is freely available at https://github.com/kyleabeauchamp/EnsemblePaper.

The regularization strength λ weights simulation versus experimental data. To determine this weighting in an unbiased manner, BELT carries out cross validation on the simulation data, as described in Appx. S5; this procedure also reduces errors due to finite sampling of equilibrium properties. For each model, we used PyMC to sample at least 5,000,000 values of *α*; sampled values of *α* were thinned 100-fold to reduce correlation. The first 5,000 samples (before thinning) were discarded as burn-in. Convergence of MCMC sampling was assessed by visual examination of MCMC traces; a well-sampled and thinned trace will appear to be white noise, without correlation between one sample and the next. MCMC traces are shown in Fig. S2 and discussed in Appx. S7. To incorporate simulation uncertainty, we used Bayesian Bootstrapping (Appx. S6). Two Bayesian bootstrap replicates were performed.

## Results

Short peptides provide crucial tests for evaluating and optimizing molecular dynamics force fields (9, 11, 14, 19, 40). Such peptides offer a window into the intrinsic conformational propensities of amino acids, free from the secondary structure bias found in statistical surveys of protein structures (41). To test the proposed theoretical framework, we used BELT to infer the conformational populations of trialanine from chemical shift and scalar coupling measurements (19).

### Conformational Propensities of Trialanine Simulations

Trialanine was simulated (see Methods) in five different force fields. The chosen force fields show considerable variation in their predicted conformational propensities. The ff96 force field shows a bias towards *β* conformations (population: 51%) (Fig. 2b, red). On the other hand, ff99 strongly favors helical conformations, with a predicted *α* population of 80% (Fig. 2c, red). The *PP*_*II*_ state, known to be the dominant state in solution from independent approaches (19, 40, 42), is the dominant simulated state only in the ff99sbnmr-ildn force field (Fig. 2a, red). Low *PP*_*II*_ populations and inconsistency between force fields have been previously noted (9, 11, 14, 19).

**Figure 2.**
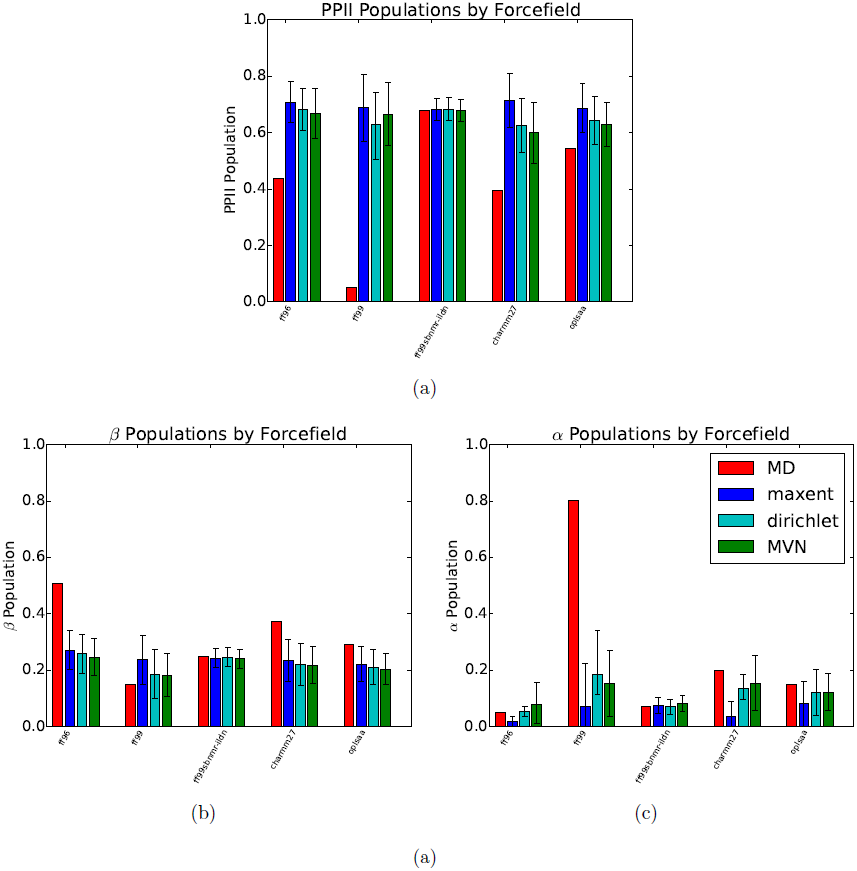
MD and BELT (maxent, Dirichlet, and MVN priors) conformational propensities (for central alanine residue) in each force field.

### Agreement with NMR Measurements: MD and BELT Ensembles

Given the differences in conformational propensities, one might expect varying degrees of agreement with the available experimental measurements. This is indeed the case; four out of five force fields show values of the reduced 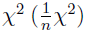 greater than 1.0 (Fig. 3a, red). Because of this considerable error, we therefore examined BELT hyperensembles based on incorporating six NMR measurements of chemicals shifts (*C*^*α*^, *C*^*β*^, and *H*) and scalar couplings (^3^*J*(*H*^*N*^*C*′), ^2^*J*(*NC*^*α*^), and ^3^*J*(*H*^*N*^*C*^*β*^)) to reweight each of the five molecular dynamics ensembles. As expected, the BELT hyperensembles accurately recapitulate these six measurements used in the reweighting (Fig. 3a). In a more incisive test, the BELT hyperensembles accurately predicted four measurements (*H*^*α*^ chemical shift and ^3^*J*(*H*^*N*^*H*^*α*^), ^3^*J*(*H*^*α*^*C*′) and ^1^*J*(*NC*^*α*^) scalar couplings) that were not used to fit the models. (Fig. 3b). A table of predicted and observed NMR measurements is given in Tables 1, S1, and S2.

**Figure 3.**
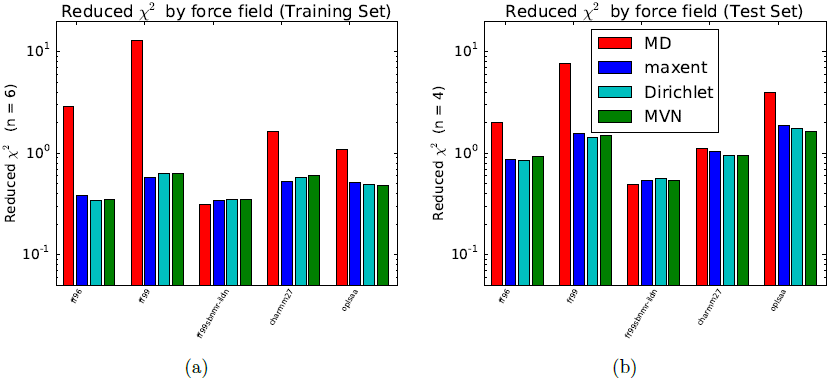
The reduced *χ*^2^ error (e.g. 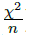) for MD and BELT (maxent, Dirichlet, and MVN priors) models. The BELT reduced *χ*^2^ is estimated as the mean reduced *χ*^2^ over all MCMC samples, (a). Calculated using the six measurements used to fit the BELT model, (b). Calculated using four measurements not used to fit the BELT model. See Methods for the definition of training and test sets. Note that the training and test sets are not fully independent because all measurements probe the (*ϕ*, *ψ*) backbone torsions.

**Table 1.** Predicted and measured observables are given. BELT predictions are calculated using the maxent prior; see Tables S1-S7 for complete table. The ‘all‘, ‘training‘, and ‘test‘ datasets have 10, 6, and 4 measurements, respectively.

| | $F_i$ | $\sigma_i$ | ff96 | | ff99 | | ff99sbnmr-ildn | | CHARMM27 | | OPLS-AA | |
| --- | --- | --- | --- | --- | --- | --- | --- | --- | --- | --- | --- | --- |
|  |  |  | MD | BELT | MD | BELT | MD | BELT | MD | BELT | MD | BELT |
| $C^\alpha$ | 52.4 | 0.9 | 52.1 | 52.2 | 52.8 | 52.5 | 52.4 | 52.4 | 52.5 | 52.4 | 52.2 | 52.2 |
| $C^\beta$ | 19.2 | 1.0 | 20.0 | 19.8 | 18.0 | 18.8 | 18.3 | 18.4 | 18.2 | 18.6 | 19.6 | 19.6 |
| $H$ | 8.6 | 0.5 | 8.6 | 8.6 | 8.3 | 8.4 | 8.2 | 8.2 | 8.3 | 8.3 | 8.6 | 8.6 |
| $H^\alpha$ | 4.4 | 0.2 | 4.6 | 4.6 | 4.6 | 4.6 | 4.5 | 4.5 | 4.6 | 4.6 | 4.6 | 4.6 |
| $^1J(NC^\alpha)$ | 11.3 | 0.5 | 11.3 | 11.5 | 10.4 | 11.8 | 11.5 | 11.5 | 11.2 | 11.7 | 11.1 | 11.3 |
| $^3J(H^\alpha C\iota)$ | 1.8 | 0.4 | 2.0 | 1.7 | 2.2 | 1.7 | 1.8 | 1.8 | 2.0 | 1.8 | 2.2 | 2.0 |
| $^3J(H^N C^\beta)$ | 2.4 | 0.2 | 1.5 | 2.3 | 0.8 | 2.3 | 2.3 | 2.3 | 1.8 | 2.3 | 1.9 | 2.2 |
| $^3J(H^N C\iota)$ | 1.1 | 0.3 | 1.5 | 1.2 | 1.8 | 1.2 | 1.0 | 1.0 | 1.4 | 1.2 | 0.9 | 0.8 |
| $^3J(H^N H^\alpha)$ | 5.7 | 0.4 | 6.6 | 5.7 | 7.5 | 5.6 | 6.1 | 6.0 | 6.3 | 5.7 | 7.0 | 6.5 |
| $^2J(NC^\alpha)$ | 8.4 | 0.5 | 8.5 | 8.6 | 6.4 | 8.5 | 8.5 | 8.5 | 8.1 | 8.6 | 8.1 | 8.4 |
| $\chi^2$ (all) | | | 2.5 | 0.6 | 10.8 | 1.0 | 0.4 | 0.4 | 1.4 | 0.7 | 2.3 | 1.1 |
| $\chi^2$ (train) | | | 2.9 | 0.4 | 12.9 | 0.6 | 0.3 | 0.3 | 1.6 | 0.5 | 1.1 | 0.5 |
| $\chi^2$ (test) | | | 2.0 | 0.9 | 7.6 | 1.6 | 0.5 | 0.5 | 1.1 | 1.0 | 4.0 | 1.9 |

### Converged Conformational Propensities Observed in BELT Ensembles

Although the raw MD simulations predicted quite different conformational propensities, BELT reweighting gave five ensembles with conformational populations that agreed to within estimated statistical uncertainty (Fig. 2). Quantitative predictions and uncertainties are given in Tables S3-S6. In accord with expectation, the lower accuracy force fields (e.g., ff99) were assigned lower λ values than force fields that were able to predict the experimental data *a priori* (see Supporting Information). The lower accuracy simulations also give final predictions that were more uncertain (e.g., PP_II_ frequency of 69 ± 13% for ff99) than force fields that are able to predict experimental data *a priori* (e.g., *PP*_*II*_ frequency of 71 ± 4% for ff99sbnmr-ildn). Nevertheless, the final predictions agreed, and residual modest differences provided practical estimates of systematic error. In general, we find (*PP*_*II*_, *β*, *α*) populations of (67 ± 9%, 23± 6%, 10± 8%); here the mean and uncertainty are approximated as the mean and standard deviation across all force fields and priors.

In addition to convergence between models constructed from different force fields, we also assessed the convergence between BELT models built using different priors on the parameters *α*. In general, different priors gave similar results with small quantitative differences (Figs. 2 and 3). Building BELT models with different priors could therefore be further useful for bracketing uncertainties in situations with limited simulation data.

### The Resolution Limit of Trialanine BELT Ensembles

Despite the near-quantitative agreement in *α*, *β*, and *PP*_*II*_ populations (Fig. 2) and overall Ramachandran features (Fig. 4), the fine details of the Ramachandran plots differed between the five models. Because all five BELT ensembles showed excellent agreement with experiment (Fig. 3), we concluded that six chemical shifts and scalar couplings were insufficiently informative to resolve (and falsify) subtle force field differences. The most obvious such difference was the width, shape, and orientation of the *PP*_*II*_ basin. Most strikingly, ff96 and OPLS-AA gave *PP*_*II*_ basins that were vertically oriented in the Ramachandran plot, while ff99, ff99sbnmr-ildn, and CHARMM27 gave diagonally oriented *PP*_*II*_ basins. Two different effects contributed to this resolution limit: the information content in the experimental measurement and the uncertainty in predictors of experimental observables. Again, the BELT strategy of modeling with different starting MD simulations revealed the residual uncertainties from these systematic errors.

**Figure 4.**
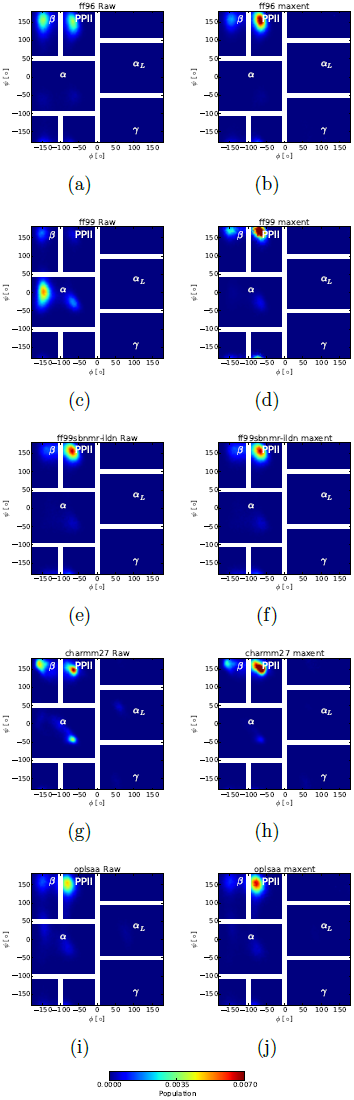
Ramachandran plots illustrate discordance of raw MD ensembles and final agreement of BELT (maxent prior) ensembles over the five tested force fields. Results from alternative BELT priors are shown in Fig. S1. The jagged appearance of the ff99 BELT model is due to limited sampling of PPII configurations in that forcefield.

## Discussion

### Structural Ensemble Biology

Why model structural ensembles, rather than just structures? At least three compelling reasons favor ensembles. First, biological molecules are multi-state machines that fold, unfold, bind ligands, aggregate, and change conformation. Biology is controlled by the relative populations of these states. Ensembles capture aspects of these phenomena by encoding equilibrium populations with structures. A second argument for ensemble modeling is fidelity to experiment. Most solution experiments measure ensemble average equilibrium properties: chemical shifts, scalar couplings, NOEs, SAXS, and FRET can often be approximated as equilibrium properties. A truly quantitative connection to these measurements requires modeling the equilibrium ensemble. Finally, recent advances in atomistic simulation (34, 43, 44, 45), special-purpose hardware (46), and distributed computing analysis (47, 48) have enabled atomistic simulations to reach the millisecond timescale (49, 50, 51, 52); the computational cost of ensemble modeling is quickly becoming manageable.

One might argue that structural ensembles are unnecessary because many proteins occupy a single state under physiological conditions. For such proteins, it is probably safe to enforce single state behavior, as is assumed in current modeling approaches. However, we suggest that the number of states be inferred—not assumed.

### Comparison to Previous Ensemble Methods

Previous ensemble modeling efforts that are most similar to BELT share three key ingredients: state decomposition, a *χ*^2^ objective function, and population inference on the clusters. For example, this general recipe describes the approach used in previous analyses of homopeptides (19), the EROS technique for SAXS modeling (18), and the Bayesian Weighting (BW) formalism (24, 53). Note that of these three techniques, only BW goes beyond returning a single best-fit ensemble and instead characterizes the posterior distribution via MCMC; below we therefore focus our attention on BW as it is most directly comparable to BELT in scope and purpose.

The primary disadvantage of previous techniques is the need for a state decomposition, which must be defined either by hand or by clustering. Working with a given state decomposition can introduce two different errors, depending on the number and quality of states. In the limit of few states, clustering can overly coarsen the system of interest, preventing the model from reproducing multiple experimental observables. At the other extreme, too many states leads to a large number of parameters to be estimated. This will lead to poor generalization performance and large errors when predicting experiments not used to train the model, as well as reliance on a subjective choice of how many states is appropriate.

One symptom of this regime is discontinuity in conformational populations. For example, imagine two nearby conformations at the boundary between two BW states—one conformation on each side of the boundary. In BW, the populations of each conformation could fluctuate dramatically with the corresponding state populations. In BELT, however, the two conformations will have nearly identical populations if the predicted observables vary smoothly.

BELT avoids arbitrary state decompositions by projecting simulations onto a basis defined only by the information at hand: the unweighted simulation and the function that maps ensembles onto observables. The advantage of working in this basis are threefold. First, in BELT, one estimates a single parameter (*α*_*i*_) for each experimental observable. If the number of experiments is small, as is often the case, the inference problem involves only a few parameters. Second, the predicted observables are a natural basis for biophysical calculations, in that the predicted observables are the fundamental connection between simulation and experiment. Working in this basis allows direct connection to experiment and often provides insight into the molecular interactions driving biophysical phenomena. For example, the projection onto observables could be used to rationally infer force field parameters—essentially a Bayesian version of the ForceBalance method (54, 55). Third, BELT does not require subjective choices. In the limit of exact measurements, BELT reduces to a previous (15) maximum entropy approach, and, more generally, is connected to the Crooks hyperensemble formalism (see Appx. S1).

We also point out some surprising differences between BELT and BW-like methods. BW- like methods have the property that the in-state means of features are preserved, leading to an undesirable dependence on the choice of state decomposition. More precisely, suppose that *χ*_*s*_(*x*) is the indicator function of a conformational state *s*. Then in-state averages of the form < *χ*_*s*_(*x*) >^-1^< *h*(*x*)*χ*_*s*_(*x*) > do not depend on the reweighted populations. BELT, however, does not preserve the in-state averages; in fact, this property is the direct result of BELT’s connection to maximum entropy modeling (see Appx. S1 and ref. (15)). The effect of this property is that the peaks of reweighted histograms are slightly shifted relative to the raw MD results, as observed in Fig. S4. We conclude that BW-like methods are useful for systems with few, well-defined conformational states, while BELT may offer significant advantages in the absence of an obvious state decomposition.

In addition to BW-like methods, there are also a class of methods where restrained simulations are used to derive ensembles of hundreds of conformations that, when taken together, produce the correct ensemble average observables (5, 56). Through the use of restraints, such methods have advantages in situations where the initial force field is insufficiently accurate to sample the correct regions of conformation space. Unlike BELT, however, these methods do not yet give a statistical treatment of uncertainty from errors in experiments or connecting simulations to experiments; new predictions cannot be rigorously falsified or validated in subsequent experiments.

More recently, a similar Bayesian technique for structural ensemble inference has been developed (57).

### Comparison to a Previous Trialanine Study

Our results are in qualitative, but not quantitative, agreement with a previous study of trialanine (19) using the same experimental measurements. That study suggested a *PP*_*II*_ population as high as 92 ± 5%, somewhat higher than our 67 ± 9% and with a twofold lower estimated uncertainty. The difference can be attributed to three methodological differences. First, the previous study used likelihood maximization to directly fit the (*PP*_*II*_, *β*, and *α*) populations from a three-state decomposition of their simulations. The use of likelihood maximization may give misleading results when the likelihood surface is broad and shallowly peaked, as was found in the previous study. However, this does not appear to be the primary cause of disagreement, as maximization of the BELT likelihood recovers populations within ±5% of the values obtained via MCMC sampling. Second, the previous study assumed each scalar coupling to have an uncertainty of 1, while we approximate the uncertainties as the RMS errors determined when fitting the Karplus equations. This weights the measurements differently and will lead to quantitative differences in estimated populations. Different choices of Karplus coefficients also may lead to different predicted properties, as has been discussed elsewhere (9, 58). Finally, the prior method’s choice of state decomposition may cause slight differences in estimated conformational populations.

### Performance and Extension to Larger Systems

The computational performance of BELT depends on several factors. First and foremost, the cost is proportional to the number of requested MCMC samples (*n*_*samples*_). The required number of samples must be determined by convergence analysis of the resulting MCMC traces. Second, each step of MCMC sampling requires calculations involving each member of the *m* conformations in the ensemble; *m* is the second major determinant of computational cost. Finally, each step of MCMC sampling involves drawing one random variable for each of the *n* experimental measurements, so the cost of each MCMC step depends (albeit weakly) on *n*. For the present work (*m* = 3 × 10^5^, *n* = 6, *n*_samples_ = 10^7^), each BELT run required approximately 2 days of compute time on an Intel 3770K processor. A similar calculation with only a single experimental observable (*n* = 1) would take approximately 1.8 days. For a larger system, say ubiquitin with one measurement per residue, one might work with fewer conformations to reduce the computational cost. As an example of the computational cost, a calculation with (*m* = 5 × 10^4^, *n* = 76, *n*_samples_ = 10^7^) would require approximately 1 day.

Because the present analysis has focused on the analysis of a small peptide, we briefly discuss two possible challenges in applying BELT to larger protein systems. First, the computational cost of molecular dynamics simulations currently prevents converged equilibrium simulations of full protein systems; this was one motivation for our choice of trialanine as a model system. Second, inaccurate force fields may reduce the overlap between the true ensemble and that sampled in simulation. Given a finite simulation length, it is possible that no amount of reweighting could provide agreement with experiment. Force field inaccuracy may become increasingly important for larger protein systems (59).

## Conclusion

Bayesian Energy Landscape Tilting allows the simultaneous characterization of structural and equilibrium properties by generating a Bayesian ensemble of conformational ensembles— a hyperensemble. Through its use of MCMC, BELT is robust to ambiguous experiments and provides rigorous uncertainty estimates, as illustrated here in the case of a tripeptide system with a complex ensemble. BELT models constructed with a handful of NMR measurements correct significant force field bias, provide generalizable, force field independent trialanine ensembles, and allow evaluation of residual systematic errors. Important frontiers for BELT include the integration of numerous rather than sparse data and extension of the current equilibrium framework to prediction of kinetic properties. The principled combination of simulation and experiment—and evaluation of convergence from multiple force fields—will enable predictive models that might not be achievable using either simulation or experiment alone.

## Acknowledgements

We thank John Chodera, TJ Lane, Frank Cochran, Pehr Harbury, Xuesong Shi, and Dan Herschlag for helpful discussions.

## Supporting Information

Tables S1-S7, Figures S1-S4, and Appendices S1-S7 are available in the supporting information.

## Supporting References

References 60-62 appear in the supporting information.

## References

[1] Berman, H. M., J. Westbrook, Z. Feng, G. Gilliland, T. N. Bhat, H. Weissig, I. N. Shindyalov, and P. E. Bourne, 2000. The Protein Data Bank. Nucleic Acids Res. 28:235–235.

[2] Dethoff, E. A., K. Petzold, J. Chugh, A. Casiano-Negroni, and H. M. Al-Hashimi, 2012. Visualizing transient low-populated structures of rna. Nature.

[3] Fink, A. L., 2005. Natively unfolded proteins. Curr. Opin. Struct. Biol. 15:35–35.

[4] Korzhnev, D. M., X. Salvatella, M. Vendruscolo, A. A. Di Nardo, A. R. Davidson, C. M. Dobson, and L. E. Kay, 2004. Low-populated folding intermediates of Fyn SH3 characterized by relaxation dispersion NMR. Nature 430:586–586.

[5] Lindorff-Larsen, K.., R. B. Best, M. A. DePristo, C. M. Dobson, and M. Vendruscolo, 2005. Simultaneous determination of protein structure and dynamics. Nature 433:128–132.

[6] Lange, O. F., N.-A. Lakomek, C. Fares, G. F. Schroder, K. F. Walter, S. Becker, J. Meiler, H. Grubmuller, C. Griesinger, and B. L. de Groot, 2008. Recognition dynamics up to microseconds revealed from an rdc-derived ubiquitin ensemble in solution. Science 320:1471–1471.

[7] DePristo, M. A., P. I. de Bakker, and T. L. Blundell, 2004. Heterogeneity and inaccuracy in protein structures solved by x-ray crystallography. Structure 12:831–831.

[8] Lang, P. T., H.-L. Ng, J. S. Fraser, J. E. Corn, N. Echols, M. Sales, J. M. Holton, and T. Alber, 2010. Automated electron-density sampling reveals widespread conformational polymorphism in proteins. Protein Science 19:1420–1420.

[9] Best, R., N. Buchete, and G. Hummer, 2008. Are current molecular dynamics force fields too helical? Biophys. J. 95:L07–L09.

[10] Lindorff-Larsen, K.., P. Maragakis, S. Piana, M. Eastwood, R. Dror, and D. Shaw, 2012. Systematic validation of protein force fields against experimental data. PloS one 7:e32131.

[11] Beauchamp, K., Y. Lin, R. Das, and V. Pande, 2012. Are protein force fields getting better? a systematic benchmark on 524 diverse NMR measurements. J. Chem. Theory Comput. 8:1409.

[12] Li, D.-W., and R. Bruschweiler, 2011. Iterative optimization of molecular mechanics force fields from NMR data of full-length proteins. J. Chem. Theory Comput. 7:1773.

[13] Best, R. B., X. Zhu, J. Shim, P. E. Lopes, J. Mittal, M. Feig, and A. D. MacKerell, 2012. Optimization of the additive charmm all-atom protein force field targeting improved sampling of the backbone φ, ψ and side-chain χ_1_ and χ_2_ dihedral angles. J. Chem. Theory Comput..

[14] Nerenberg, P., and T. Head-Gordon, 2011. Optimizing protein-solvent force fields to reproduce intrinsic conformational preferences of model peptides. J. Chem. Theory Comput. 7:1220–1220. http://pubs.acs.org/doi/abs/10.1021/ct2000183.

[15] Pitera, J., and J. Chodera, 2012. On the use of experimental observations to bias simulated ensembles. J. Chem. Theory Comput. 8:3445–3445.

[16] Patil, A., D. Huard, and C. J. Fonnesbeck, 2010. Pymc: Bayesian stochastic modelling in python. Journal of statistical software 35:1.

[17] Crooks, G. E., 2007. Beyond boltzmann-gibbs statistics: Maximum entropy hyper-ensembles out of equilibrium. Physical Review E 75:041119.

[18] Rozycki, B., Y. C. Kim, and G. Hummer, 2011. Saxs ensemble refinement of escrt-iii chmp3 conformational transitions. Structure 19:109–109.

[19] Graf, J., P. Nguyen, G. Stock, and H. Schwalbe, 2007. Structure and dynamics of the homologous series of alanine peptides: a joint molecular dynamics/nmr study. J. Am. Chem. Soc. 129:1179–1179.

[20] Vogeli, B., J. Ying, A. Grishaev, and A. Bax, 2007. Limits on variations in protein backbone dynamics from precise measurements of scalar couplings. J. Am. Chem. Soc. 129:9377–9377.

[21] Rieping, W., M. Habeck, and M. Nilges, 2005. Inferential structure determination. Science 309:303–303.

[22] Friedman, J., T. Hastie, and R. Tibshirani, 2001. The elements of statistical learning, volume 1. Springer Series in Statistics.

[23] Wikipedia, 2004. Kullback-leibler divergence—Wikipedia, the free encyclopedia. http://en.wikipedia.org/wiki/Kullback%E2%80%93Leibler_divergence, [Online; accessed 15-July-2013].

[24] Fisher, C. K., A. Huang, and C. M. Stultz, 2010. Modeling intrinsically disordered proteins with bayesian statistics. J. Am. Chem. Soc. 132:14919.

[25] Fisher, C. K., and C. M. Stultz, 2011. Constructing ensembles for intrinsically disordered proteins. Current opinion in structural biology 21:426–426.

[26] Rubin, D., 1981. The bayesian bootstrap. The annals of statistics 9:130–130.

[27] Kollman, P., 1996. Advances and continuing challenges in achieving realistic and predictive simulations of the properties of organic and biological molecules. Acc. Chem. Res. 29:461–461.

[28] Wang, J., P. Cieplak, and P. Kollman, 2000. How well does a restrained electrostatic potential(resp) model perform in calculating conformational energies of organic and biological molecules? J. Comput. Chem. 21:1049–1049.

[29] Li, D., and R. Bruschweiler, 2010. Nmr-based protein potentials. Angew. Chem. 122:6930–6930.

[30] Lindorff-Larsen, K.., S. Piana, K. Palmo, P. Maragakis, J. Klepeis, R. Dror, and D. Shaw, 2010. Improved side-chain torsion potentials for the amber ff99sb protein force field. Proteins: Struct., Funct., Bioinf. 78:1950–1950.

[31] Mackerell Jr, A., M. Feig, and C. Brooks III, 2004. Extending the treatment of backbone energetics in protein force fields: Limitations of gas-phase quantum mechanics in reproducing protein conformational distributions in molecular dynamics simulations. J. Comput. Chem. 25:1400–1400.

[32] Bjelkmar, P., P. Larsson, M. Cuendet, B. Hess, and E. Lindahl, 2010. Implementation of the charmm force field in gromacs: Analysis of protein stability effects from correction maps, virtual interaction sites, and water models. J. Chem. Theory Comput. 6:459–459.

[33] Kaminski, G., R. Friesner, J. Tirado-Rives, and W. Jorgensen, 2001. Evaluation and reparametrization of the opls-aa force field for proteins via comparison with accurate quantum chemical calculations on peptides. J. Phys. Chem. B 105:6474–6474.

[34] Hess, B., C. Kutzner, D. Van Der Spoel, and E. Lindahl, 2008. Gromacs 4: Algorithms for highly efficient, load-balanced, and scalable molecular simulation. J. Chem. Theory Comput. 4:435–435.

[35] Horn, H., W. Swope, J. Pitera, J. Madura, T. Dick, G. Hura, and T. Head-Gordon, 2004. Development of an improved four-site water model for biomolecular simulations: Tip4p-ew. J. Chem. Phys. 120:9665.

[36] Han, B., Y. Liu, S. Ginzinger, and D. Wishart, 2011. Shiftx2: significantly improved protein chemical shift prediction. J. Biomol. NMR 1–15.

[37] Shen, Y., and A. Bax, 2010. Sparta+: a modest improvement in empirical NMR chemical shift prediction by means of an artificial neural network. J. Biomol. NMR 48:13–13.

[38] Li, D.-W., and R. Brüschweiler, 2012. Ppm: a side-chain and backbone chemical shift predictor for the assessment of protein conformational ensembles. J. Biol. NMR 1–9.

[39] Schmidt, J., M. Blumel, F. Lohr, and H. Ruterjans, 1999. Self-consistent 3j coupling analysis for the joint calibration of karplus coefficients and evaluation of torsion angles. J. Biomol. NMR 14:1–1.

[40] Grdadolnik, J., V. Mohacek-Grosev, R. Baldwin, and F. Avbelj, 2011. Populations of the three major backbone conformations in 19 amino acid dipeptides. Proc. Natl. Acad. Sci. U. S. A. 108:1794.

[41] Jha, A. K., A. Colubri, M. H. Zaman, S. Koide, T. R. Sosnick, and K. F. Freed, 2005. Helix, sheet, and polyproline II frequencies and strong nearest neighbor effects in a restricted coil library. Biochemistry 44:9691–9691.

[42] Avbelj, F., S. Grdadolnik, J. Grdadolnik, and R. Baldwin, 2006. Intrinsic backbone preferences are fully present in blocked amino acids. Proc. Natl. Acad. Sci. U. S. A. 103:1272.

[43] Pronk, S., S. Pall, R. Schulz, P. Larsson, P. Bjelkmar, R. Apostolov, M. R. Shirts, J. C. Smith, P. M. Kasson, and D. van der Spoel, 2013. Gromacs 4.5: a high-throughput and highly parallel open source molecular simulation toolkit. Bioinformatics.

[44] Eastman, P., M. S. Friedrichs, J. D. Chodera, R. J. Radmer, C. M. Bruns, J. P. Ku, K. A. Beauchamp, T. J. Lane, L.-P. Wang, D. Shukla, and V. S. Pande, 2012. Openmm 4: A reusable, extensible, hardware independent library for high performance molecular simulation. J. Chem. Theory Comput. 9:461–461.

[45] Eastman, P., and V. Pande, 2010. Openmm: a hardware-independent framework for molecular simulations. Comp. in Sci. Eng. 12:34–34.

[46] Shaw, D., M. Deneroff, R. Dror, J. Kuskin, R. Larson, J. Salmon, C. Young, B. Batson, K. Bowers, and J. Chao, 2008. Anton, a special-purpose machine for molecular dynamics simulation. Commun. ACM 51:91–91.

[47] Senne, M., B. Trendelkamp-Schroer, A. S. J. S. Mey, C. Schutte, and F. Noe, 2012. Emma - a software package for markov model building and analysis. J. Chem. Theory Comput. 8:2223–2223.

[48] Beauchamp, K., G. Bowman, T. Lane, L. Maibaum, I. Haque, and V. Pande, 2011. Msmbuilder2: Modeling conformational dynamics at the picosecond to millisecond scale. J. Chem. Theory Comput. 7:3412–3412.

[49] Voelz, V., G. Bowman, K. Beauchamp, and V. Pande, 2010. Molecular simulation of ab initio protein folding for a millisecond folder NTL9 (1-39). J. Am. Chem. Soc. 132:1526–1526.

[50] Bowman, G. R., V. A. Voelz, and V. S. Pande, 2011. Atomistic folding simulations of the five helix bundle protein λ6-85. J. Am. Chem. Soc. 133:664.

[51] Shaw, D. E., P. Maragakis, K. Lindorff-Larsen, S. Piana, R. O. Dror, M. P. Eastwood, J. A. Bank, J. M. Jumper, J. K. Salmon, Y. Shan, and W. Wriggers, 2010. Atomic-level characterization of the structural dynamics of proteins. Science 330:341–341.

[52] Lindorff-Larsen, K.., S. Piana, R. Dror, and D. Shaw, 2011. How fast-folding proteins fold. Science 334:517–517.

[53] Fisher, C. K., O. Ullman, and C. M. Stultz, 2012. Efficient construction of disordered protein ensembles in a bayesian framework with optimal selection of conformations. Pacific Symposium on Biocomputing. 82.

[54] Wang, L.-P., J. Chen, and T. Van Voorhis, 2012. Systematic parametrization of polarizable force fields from quantum chemistry data. J. Chem. Theory Comput. 9:452–452.

[55] Wang, L.-P., T. L. Head-Gordon, J. W. Ponder, P. Ren, J. D. Chodera, P. K. Eastman, T. J. Martinez, and V. S. Pande, 2013. Systematic improvement of a classical molecular model of water. J. Phys. Chem. B 117:9956–9956.

[56] Richter, B., J. Gsponer, P. Varnai, X. Salvatella, and M. Vendruscolo, 2007. The mumo (minimal under-restraining minimal over-restraining) method for the determination of native state ensembles of proteins. Journal of biomolecular NMR 37:117–117.

[57] Olsson, S., J. Frellsen, W. Boomsma, K. V. Mardia, and T. Hamelryck, 2013. Inference of structure ensembles of flexible biomolecules from sparse, averaged data. PloS one 8:e79439.

[58] Markwick, P. R., S. A. Showalter, G. Bouvignies, R. Brüschweiler, and M. Blackledge, 2009. Structural dynamics of protein backbone φ angles: extended molecular dynamics simulations versus experimental 3 j scalar couplings. Journal of biomolecular NMR 45:17–17.

[59] Raval, A., S. Piana, M. P. Eastwood, R. O. Dror, and D. E. Shaw, 2012. Refinement of protein structure homology models via long, all-atom molecular dynamics simulations. Proteins: Structure, Function, and Bioinformatics 80:2071–2071.

[60] Jaynes, E. T., 1957. Information theory and statistical mechanics. Physical review 106:620.

[61] Flyvbjerg, H., and H. G. Petersen, 1989. Error estimates on averages of correlated data. J. Chem. Phys. 91:461.

[62] Shirts, M., and J. Chodera, 2008. Statistically optimal analysis of samples from multiple equilibrium states. J. Chem. Phys. 129:124105.

